# MICRON learns outcome-associated representations of spatial immune microenvironments

**DOI:** 10.64898/2026.04.14.718488

**Authors:** Chi-Jane Chen, Betsy George, Luvna Dhawka, Baggio Evangelista, Natalie Stanley

## Abstract

Spatial imaging proteomics modalities, such as imaging mass cytometry, enable comprehensive identification of immune microenvironments driving disease outcomes. Identifying outcome-associated immune microenvironments from these data has proven to be complex, as it requires segmenting cells with complex shapes and reconciling spatial signatures across many heterogeneous samples. We present `MICRON`, a segmentation-free, fully automated multiple-instance learning based tool for automatic identification of outcome-linked immune microenvironments. `MICRON` learns representations of samples profiled with spatial imaging proteomics modalities, enabling more accurate prognostic and diagnostic prediction over existing approaches. As a case study, we show that `MICRON` generates a comprehensive importance map that reveals key outcome-associated immune microenvironments in brain cancer, uncovering coordinated cell-cell communication between astrocytes, NK cells, and macrophages linked to survival outcomes. `MICRON` is provided as open source software for broad use by clinicians and biologists at https://github.com/ChenCookie/micron.

## 1 Introduction

Spatial imaging proteomics technologies, such as imaging mass cytometry (IMC) and multiplexed ion beam imaging by time of flight (MIBI-TOF) can uncover the spatial tissue-immune microenvironments driving disease outcome or phenotypes [1– 5]. Prototypical spatial configurations between immune cells can be leveraged for better prognostic prediction, patient stratification, or treatment planning [3, 6, 7]. Currently, the majority of computational tools for detecting microenvironments require cell segmentation [8], posing challenges and limitations for cells with complex and non-spherical morphologies, such as, microglia [9]. We introduce `MICRON` (microenvironment-aware clinical representation learning for outcome prediction via multiple-instance learning) as a segmentation-free approach for sample featurization and microenvironment identification from multiplexed imaging proteomics modalities. Spatial signatures derived from `MICRON` prioritized outcome-associated immune microenvironments that can be leveraged for more accurate prognostic prediction and clinical immunophenotyping.

Between spatially resolved proteomics and transcriptomic modalities, there have been numerous statistical and machine learning based approaches proposed to identify tissue immune microenvironments enriched in particular disease outcomes or phenotypes [10–14]. The majority of these methods, such as, SpatialLDA [12], LEAPH [13], QUICHE [11], and graph-neural-network approaches [15] require segmentation of individual cells. While effective for canonical, spherically shaped immune cells, state-of-the-art segmentation approaches, such as Cellpose [16] and DeepCell [17], which have been trained on many manually annotated cells, may struggle with cells that do not have canonical spherical morphologies, such as, microglia and neurons [18].

CANVAS was recently [10] proposed as the first segmentation-free self-supervised approach for featurizing samples profiled with spatial multiplexed imaging modalities. For downstream identification of outcome-associated microenvironments, CANVAS requires clustering image regions across datasets based on their learned representations, and can be sensitive to clustering hyperparameters. Furthermore, CANVAS does not make distinctions between informative and uninformative image regions in training, limiting microenvironment interpretability and increasing noise in learned sample representations.

We developed `MICRON` to improve the identification of outcome-associated immune microenvironments, while learning representations for each sample that can robustly predict clinical outcomes or phenotypes. `MICRON` is a segmentation-free, self-supervised representation learning framework with several novel features. First, `MICRON` partitions the image into cohesive microenvironment regions, which form the building blocks of the learned representation required to compute a spatial immune signature for each sample (Fig. 1a). `MICRON` is trained to account for the variability in information content, such as microenvironment diversity, across image regions. Further, `MICRON` facilitates biological interpretation of outcome-associated microenvironments through an explainable SHAP framework [19]. The explainability component facilitates efficient identification of outcome-associated spatial immune microenvironments to generate mechanistic hypotheses about how cell-cell communication is tuned in disease or in response to a treatment. These innovative aspects of `MICRON` collectively facilitate its use in identifying meaningful immune microenvironments from large clinical immunophenotyping studies.

**Fig. 1.**
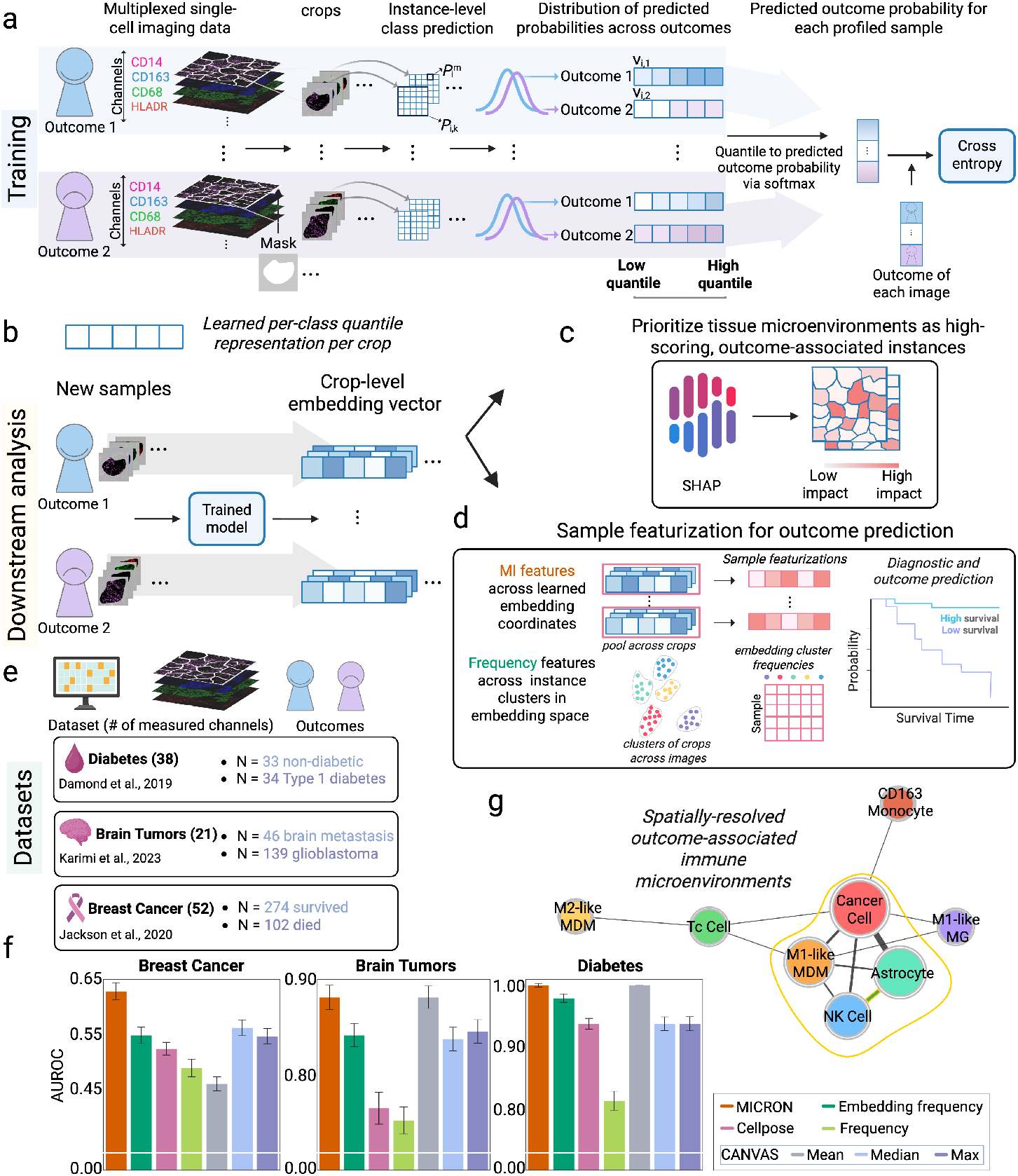
MICRON learns outcome-associated spatial immune signatures and microenvironments. (**a**) Schematic of the multiple-instance learning framework used to train `MICRON` to predict outcomes from spatial imaging proteomics modalities. During training, square regions (instances) are cropped from the input image and processed by a fully convolutional network (FCN) to generate instance-level class predictions. The distribution of instance-level outcome probability predictions are summarized via a quantile-based featurization per each outcome class. The quantile vectors across classes are concatenated and input to a softmax function to predict an outcome probability for each sample. The model parameters, including both the FCN and quantile aggregation module, are optimized via the cross-entropy loss. (**b**) Sample representations are computed for new, unseen samples by using the trained `MICRON` model to generate embedding vectors for each image crop. (**c**) Crops across testing images are prioritized via SHAP values and identify both high-impact and outcome-associated image crops, corresponding to key immune microenvironments. (**d**) For predicting sample outcomes, `MICRON` can specify two types of immune features, including (i) (multiple-instance (MI) features), obtained by pooling learned instance embedding dimensions across all cells and (ii) (frequency features), obtained by clustering instance embedding vectors and counting the proportion of instances assigned to each cluster per sample. Featurizations are used for downstream prediction tasks, such as, survival analysis. (**e**) Immune signatures obtained through `MICRON` were evaluated for their accuracy in predicting binary clinical outcomes on three publicly available clinical IMC datasets. The number of channels measured in the IMC experiments are denoted as well as the number of donors in each outcome class. (**f**) The prediction accuracy of `MICRON` is compared to related methods for predicting a binary clinical outcome across the three datasets (area under the ROC curve (AUROC)). In the Karimi dataset, `MICRON` identified more frequent co-occurrences between astrocytes, NK cells, and M1-like MDMs in glioblastoma patients than was observed in patients with metastasized brain tumors. Bold green lines indicate the most frequently co-occurring cell-type pairs across topmost-informative image crops. Nodes circled in gold represent the set of cell-types that were most ubiquitously identified across immune microenvironments.

## 2 Results

A key strength of `MICRON` is its ability to learn outcome-associated immune signatures, or composite per-sample representations, reflecting immune cell function and organization in a tissue (Fig. 1b-d). Specifically, spatial immune signatures derived through `MICRON` led to more accurate prediction of binary clinical outcomes across three IMC datasets profiling patients with diabetes [20], breast cancer [6], and brain tumors [21] (datasets and binary outcomes to be described in Fig. 1e and summarized in Methods), in comparison to those obtained by CANVAS [10] and non-learning-based approaches (outlined in Supplementary Fig. S1). `MICRON` learned informative spatial immune signatures in the breast cancer dataset, enabling significantly higher classification of patients according to their survival outcome (Fig. 1f). Specifically, a model trained with `MICRON` predicted survival outcome in a held-out cohort of patients with (AUC = 0.63), which was significantly higher than the accuracy obtained with CANVAS (median pooling) as the second-highest performer (AUC = 0.56). `MICRON` similarly showed strong performance in classifying patients with metastatic brain cancer from glioblastoma (AUC = 0.88) in line with the accuracy obtained by CANVAS (mean pooling) (AUC = 0.88). However, `MICRON` also complements this strong classification accuracy by identifying outcome-associated microenvironments through co-occurrence patterns between cell-types from SHAP-prioritized image crops [19] (Fig. 1c-d). In the brain tumors dataset, we identified prominent spatial co-occurrences between astrocyte–natural killer (NK) cell, astrocyte–M1-like monocyte derived macrophages (MDM), and M1-like MDM–NK cells in patients with glioblastoma (Fig. 1g). Finally, `MICRON` and CANVAS (mean pooling) showed superior performance (both achieving AUC = 1.0) in classifying non-diabetic and type 1 diabetes patients, and was closely followed by the accuracy obtained using frequencies computed on the superpixel-level embedding vectors computed across the images (AUC = 0.97). Our results suggest that `MICRON` offers robust classification across diverse clinical datasets, without any segmentation, while also prioritizing outcome-associated cellular microenvironments. We hypothesize that `MICRON` achieves higher accuracy in the breast cancer dataset because it can better identify specialized, prognostic niches that may be obscured due to the disease heterogeneity.

`MICRON` was trained to classify patients with brain metastasis (BrM) from glioblastoma (GBM) patients, based on 127 multiplexed training images from the Karimi study [21]. In doing so, `MICRON` successfully prioritized brain microenvironments across 46 test set images, which were distinct between BrM and GBM patients. Fig. 2a shows representative images for a BrM patient and a GBM patient, colored by markers used to distinguish cell-nuclei (DNA1), macrophages (CD68), astrocytes (GFAP), and NK cells (CD94/CD16). Zooming in on the top two highest scoring image regions (under `MICRON`) for distinguishing BrM from GBM patients in a set of 46 held-out patients, we observed that astrocytes and NK cells tended to be more spatially proximal in GBM patients than in BrM patients (Fig. 2a-middle-right; additional details for cell phenotyping are shown in Fig. S3). Learned representations (embeddings) for the 20 top-most-informative superpixels (as defined in Methods) across test set images were clustered into *k* = 10 clusters, or prototypical microenvironment patterns across the 46 test set images (Fig. 2b-c). We classified each donor as having high abundance of a microenvironment (cluster) if its frequency of the microenvironment was in the top 25% across the 46 test set samples, which uncovered five statistically significant (*p* ≤ 0.001) survival-associated microenvironments (Fig. S2). Patient survival given by the Karimi study was defined as the number of days survived post-surgery, and microenvironments 4 and 9 are shown as representative survival-associated and not associated with survival microenvironments, respectively (Fig. 2c). Kaplan-Meier survival analysis revealed longer survival times associated with a high abundance of cluster 4, mediated by a coordinated spatial signature between NK cells, astrocytes, and macrophages (Fig. 2d).

**Fig. 2.**
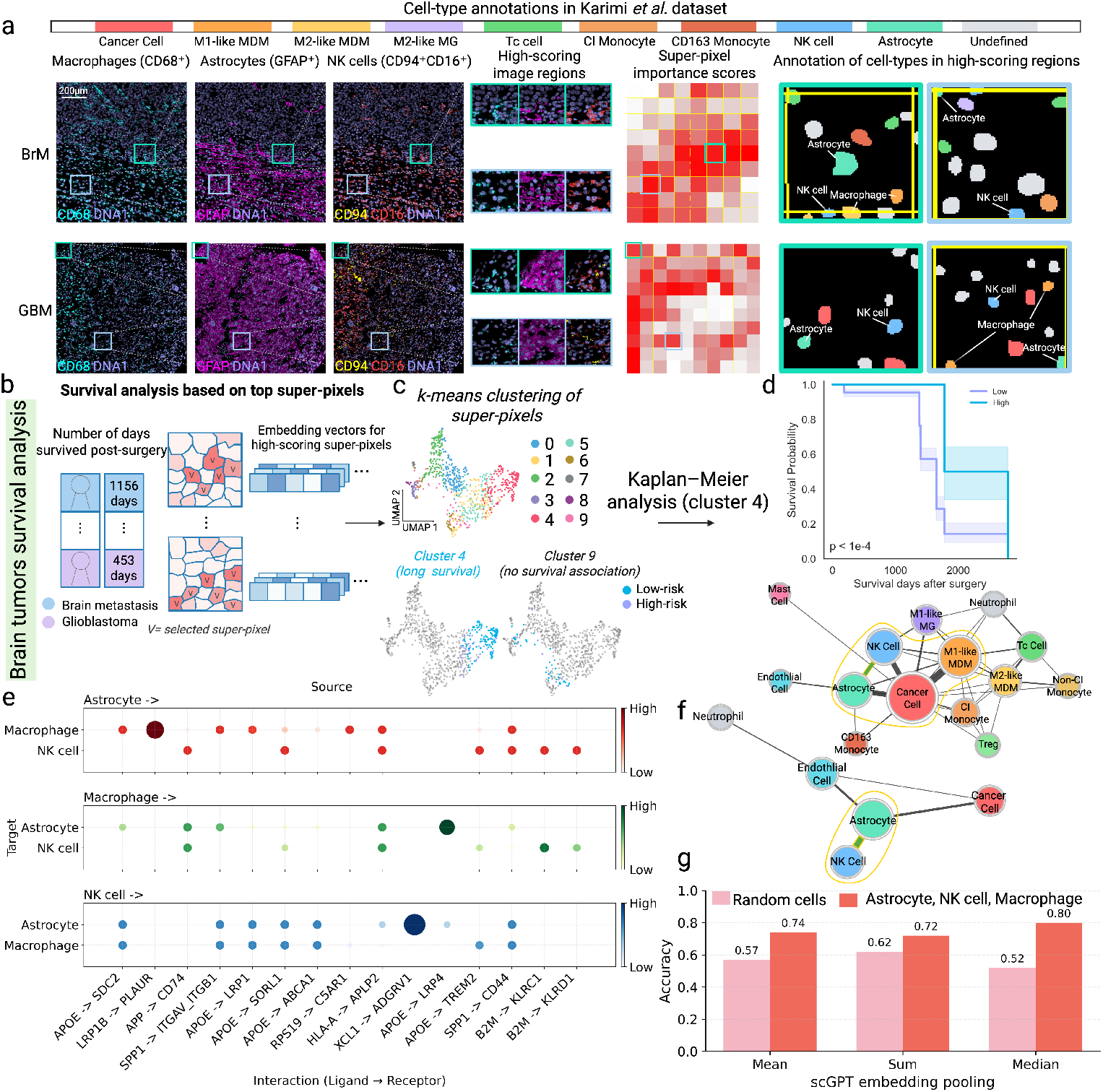
Microenvironments uncovered with MICRON prioritized differential cell-cell communication mechanisms. (**a**) Representative brain metastasis (BrM) and glioblastoma (GBM) sample images colored according to expression of key markers for phenotyping macrophages (CD68), astrocytes (GFAP), and NK cells (CD94 and CD16). Superpixel importance scores show image regions prioritized by `MICRON`. Cells in high-scoring regions were segmented and phenotyped to reveal prominent microenvironments in the BrM and GBM classes. (**b**) Overview of how `MICRON` can be applied to perform survival analysis in patients from the Karimi study. For a set of held out patients, `MICRON` was used to identify the top 20 most outcome-associated superpixels in each image (denoted as V in superpixel map). (**c**) Image crops were clustered with *k*-means (*K* = 10) using `MICRON`-computed embeddings. The UMAP (top) shows cells colored according to the embedding clusters. Patients were stratified into high and low frequency groups for each cluster. The UMAP (bottom) colors cells based on the risk of death and identifies clusters 4 and 9 as being survival-associated and survival unassociated, respectively. (**d**) Cluster 4 was observed to be strongly associated with survival and highlights strong spatial co-occurrence between astrocytes, NK cells, and M1-like MDMs in glioblastoma. (**e**) Cell-cell communication patterns via ligand/receptor signaling were investigated in an independent cohort of glioblastoma patients profiled with single-cell RNA-seq (Ruiz-Moreno study [22]). Dotplots visualize top genes with enrichment patterns according to specificity rank (indicated by dot size and color) specifically between astrocytes, macrophages, and NK cells. (**f**)A graph was constructed between cells where the edges are weighted by the specificity rank between a pair of cell-types. The graph reveals prominent cross-talk between astrocytes and NK cells, supporting the spatial signature prioritized by `MICRON` in the Karimi study. (**g**) scGPT was used to featurize samples in the Ruiz-Moreno glioblastoma study based on genes prioritized by LIANA for mediating ligand-receptor mediated signaling between astrocytes, NK cells, and macrophages. Featurizations were used to predict the age of each donor under different pooling strategies (mean, sum, and median) in the implementation of scGPT. Prediction accuracy from scGPT features (pink) was compared to that obtained using featurizations obtained using random cells and genes (pink).

The confidence of spatial signatures prioritized by `MICRON` can be further bolstered by considering ligand-receptor mediated signaling events that are enriched among transcriptomic measurements obtained in other glioblastoma studies. We evaluated the extent to which the survival-linked signature between NK cells, astrocytes, and macrophages, specifically, could be mechanistically supported across 240 glioblastoma patients profiling the brain with single-cell RNA sequencing (scRNA-seq) from the Ruiz-Moreno study [22]. Using LIANA [23], a consensus tool for identifying enriched ligand-receptor interactions, significant interactions were predicted between NK cells and astrocytes (Fig. 2f), mediated between XCL1 (ligand in NK cells) and ADGVR1 (receptor in astrocytes). This suggests a plausible mechanism for communication between these cell-types mediated through their spatial proximity that is associated with longer survival times in patients with brain cancer (Fig. 2e-f).

Spatial immune microenvironments uncovered with `MICRON`, complemented by enriched ligand-receptor signaling uncovered in transcriptomic studies provide rich signals that can be leveraged to generate robust immune signatures for diagnostic prediction. We computed scGPT embeddings [24] in the Ruiz-Moreno study [22] based on the set of genes relevant to ligand-receptor mediated cell-cell communication between astrocytes, NK cells, and macrophages that were observed to be enriched across these respective cells in the Karimi dataset. Immune signatures were computed by pooling scGPT embedding dimensions across astrocytes, NK cells, and macrophages in each sample and used to predict a binary age classification (age *<* 60 vs. ≥60) for each donor. The ability to predict age, or any clinical outcome, based on gene expression signatures in astrocytes, NK cells, and macrophages suggests that their cell-cell communication is a tunable circuit across clinical phenotypes. Median pooling on scGPT embedding coordinates resulted in strong accuracy in age prediction (accuracy = 0.8), in contrast to the best result obtained selecting random cells and genes from each donor (accuracy = 0.62 under sum pooling) (Fig. 2g). This suggests that the genes relevant to ligand-receptor based signaling between astrocytes, NK cells, and macrophages prioritized by `MICRON` can be leveraged to engineer immune signatures for robust outcome prediction, or can form mechanistic hypotheses for future studies.

## 3 Discussion

`MICRON` leverages a scalable multiple-instance learning approach for learning outcome-associated spatial immune signatures from samples profiled with multiplexed imaging proteomics modalities. The segmentation-free nature of `MICRON` facilitates its application across tissues and disease contexts, particularly with respect to cells with complex morphologies, such as, neurons and microglia [9]. `MICRON` also provides an explainable framework for identifying and characterizing the most ubiquitous and information-rich outcome-associated immune microenvironments in an automated manner. `MICRON` can be used in clinical immunophenotyping studies to generate hypotheses about spatial immune microenvironments driving disease outcomes and phenotypes. As a key contribution, `MICRON` presents a segmentation-free, self-supervised representation learning framework that effectively captures informative regions within each sample. Removing the requirement for cell segmentation prevents potential biases introduced by segmentation models and variations in image quality. Furthermore, `MICRON` focuses on learning representations from multiplexed imaging data by selectively leveraging informative superpixels, or spatially contiguous regions, rather than processing the entire image or randomly sampled regions. This strategy thereby primes the model to learn image representations driven by dense cellular regions, in contrast to potentially less-informative background information [25].

`MICRON` complements existing automated approaches [11, 13, 15], in identifying outcome-associated tissue microenvironments without segmentation. Specifically, `MICRON` is the first approach to specify a map over the entire tissue image to prioritize spatial immune signatures driving an outcome. These spatial insights can be used to prioritize ligand-receptor mediated cell-cell communication mechanisms that may mediate the spatial proximity between cells [26–28]. Specifically, we applied the methodology to understand the brain immune microenvironments that differ between primary glioblastoma and metastatic brain tumors. These analyses highlighted co-occurrence between astrocyte–NK cell, astrocyte–M1-like MDM, and M1-like MDM–NK cells. Spatially, we have also discovered the co-occurrence between astrocyte, macrophage, and NK cell in the Ruiz-Moreno study, which is an independent single-cell transcriptomic study. This aligns with previous studies describing the role of reactive astrocytes in modulating NK cell-mediated immune surveillance [29] and their involvement in recruiting monocyte-derived macrophages to the brain TME [30, 31]. Featurizations learned through `MICRON` can also stratify patients by survival, enabling rapid identification of survival-linked immune microenvironments.

This methodology can be expanded in future works by generating immune signatures for outcome prediction that can jointly take into account the spatial signatures gleaned through `MICRON` with other omics or clinical measurements [32–34]. Future work could adapt `MICRON` to learn continuous outcomes to enable mapping of how spatial signatures change across dynamic processes, such as cellular processes, disease progression, or age [35–37].

Overall, `MICRON` pioneers a new direction for inferring outcome-linked immune microenvironments from spatial multiplexed imaging proteomics data. `MICRON` facilitates biological interpretability and usability for biologists and clinicians by prioritizing key image regions and immune microenvironments linked to disease outcomes or survival patterns without requiring any segmentation. Analysis with `MICRON` may prioritize cellular interactions that can be harnessed in immunotherapy approaches.

## 4 Methods

`MICRON` is a segmentation-free machine learning approach to learn outcome-associated immune signatures from spatial multiplexed imaging proteomics data in a scalable manner. In doing so, we identify information-rich sections of the image which we interpret to be key immune microenvironments associated with an outcome of interest. Our solution leverages a multiple-instance learning framework [38–40], where small crops or subsets of the training images are used to train a model to predict a clinical label.

### 4.1 Datasets

We evaluated `MICRON` using three publicly available imaging mass cytometry (IMC) datasets profiling clinical cohorts with diabetes (Damond *et al*. [20]), breast cancer (Jackson *et al*. [6]), and brain tumors (Karimi *et al*. [21]). We also complemented spatial insights uncovered through `MICRON` with mechanistic insights obtained from a complementary single-cell RNA-sequencing dataset profiling patients with glioblastoma [22]. The details of these datasets are summarized below.

#### Imaging Mass Cytometry (IMC) Datasets

##### Diabetes dataset (Damond *et al*. [20])

The IMC dataset generated by Damond *et al*. longitudinally profiles pancreatic tissue samples obtained from 67 human donors, including both non-diabetic individuals and patients diagnosed with Type 1 diabetes (T1D). The dataset is comprised of 33 non-diabetic controls and 34 donors with established long-standing T1D. Long-standing T1D patients have a prolonged disease duration and show sustained autoimmune-mediated *β*-cell depletion and chronic inflammatory remodeling of the pancreatic microenvironments [41]. The IMC panel included 38 channels, where 35 of the channels measured protein markers. In our classification experiments, we aimed to distinguish non-diabetic donors from donors with long-standing T1D.

##### Brain tumors dataset (Karimi *et al*. [21])

The brain tumors IMC dataset [21] profiles the brain tumors microenvironment from 139 patients with primary glioblastoma (GBM) and 46 individuals with metastatic brain tumors (BrM). For each sample, we considered 18 common channels (17 protein markers) that were shared across samples. The samples were obtained across the brain tumors microenvironments, including tumor core, invasive margin, and adjacent non-malignant tissue compartments. Because certain patients contributed multiple spatial regions, the dataset includes 270 GBM samples and 123 BrM samples. In classification experiments, we sought to classify patients with glioblastoma from those with metastatic brain tumors.

##### Breast cancer dataset (Jackson *et al*. [6])

The breast cancer dataset [6] profiles primary breast tumor tissues with IMC. Associated with the IMC data were clinically annotated follow-up information, including patient age, tumor size, treatment history, and survival status. The IMC panel stained for 52 channels, where 40 of the channels measured protein markers. We note that there are multiple molecular subtypes of breast cancer, including positive or negative estrogen receptor (ER), progesterone receptor (PR), and human epidermal growth factor receptor 2 (HER2). While there are likely subtype-specific effects on the tumor-immune microenvironments, here we are only focusing on learning representations of IMC images to classify patients according to their survival outcomes (*alive* and *deceased*). Survival is an endpoint that reflects patient prognosis and captures the combined effects of tumor biology and the surrounding immune microenvironments [42]. The full dataset is comprised of 376 tumor samples, including 274 patients who were alive at last follow-up and 102 patients who had died.

#### Single-cell RNA-sequencing datasets

##### GBMap Glioblastoma Dataset (Ruiz *et al*.) [22]

The GBMap glioblastoma dataset [22] is comprised of single-cell measurements obtained from the brain tumors microenvironments from 240 patients, with associated clinical metadata, such as age, at diagnosis. We binned each patient’s age to define a binary outcome variable, classifying patients as either *young* (*<* 60 years) or *old* (≥ 60 years) groups. Previous studies identified age-dependent changes in the tumor immune microenvironments with survival [43], which we sought to link to immune signatures generated through our approach. For downstream analysis, we retained only brain cell-types that were also present in the Karimi *et al*. brain tumors dataset. This resulted in 1,042,364 cells with 26,302 genes used in all subsequent analyses.

### 4.2 Related methods used to featurize samples profiled with IMC

We compared featurizations of the cellular tissue microenvironments generated through `MICRON` to several related methods for converting multiplexed imaging data into sample-representations or *featurizations* for downstream prediction tasks. Related methods are comprised of both segmentation-free and segmentation-based approaches, as well as clustering-based frequency features and pooling-based features learned through `MICRON`. Schematic illustrations of these works are provided in (Fig. S1b).

#### MICRON-derived features

`MICRON` **embedding frequency features** `MICRON` embedding frequency features were created by first using `MICRON` to compute representations for superpixels obtained across images. Superpixels were clustered according to their learned embedding coordinates with *k*-means (*K* = 30). For each sample, we computed frequency features as the fraction of superpixels assigned to each superpixel-based embedding cluster [44]. This *K*-dimensional frequency representation was used in downstream outcome-prediction tasks subsequently used as input to a downstream classifier to predict the outcome for each sample.

#### Featurizations derived from existing segmentation and segmentation-free approaches

##### Cellpose-based pooling featurization

We developed a featurization based on Cellpose [16] segmentations of individual cells. To do so, we first selected three markers to define the nuclear and cytoplasmic compartments to run the standard Cellpose algorithm (diabetes: H3, CD99, FOXP3; brain tumors: CD3, CD68, GFAP; breast cancer: H3, CD68, CD3). Cellpose identified individual cell boundaries and estimated cell centers. For *F* measured proteins in each dataset, we defined a *F* -dimensional featurization for each sample, reflecting the median expression of each of the *F* proteins over all of the segmented cells. This featurization is akin to *pseudobulk* in standard single-cell analysis.

##### Cellpose-based frequency featurization

We used cell segmentations obtained through Cellpose to compute frequency-based featurizations for each sample. To do so, cells identified through Cellpose were clustered across all samples based on their *F* measured proteins through *k*-means (*K* = 12). For each sample, we calculated the proportion of their cells across clusters to establish a frequency-based representation, summarizing cellular composition.

##### CANVAS pooling-based featurization

We used CANVAS [10], a self-supervised machine learning approach, to learn representations for local microenvironment (LTME) tiles in each IMC-profiled image. For each sample, we applied pooling operations (mean, median, or max) across tile embeddings to obtain a featurization for each sample.

#### Featurization for single-cell RNA sequencing data (scRNA-seq)

##### scGPT pooling-based features

scGPT [24] is a transformer-based foundation model trained on large cellular atlases to learn contextualized representations for single-cell data profiled by single-cell RNA sequencing (scRNA-seq), based on their transcriptomic features. To compute sample featurizations from scGPT representations, we used mean, median, and sum pooling to aggregate the 64 scGPT dimensions across all cells in each sample.

### 4.3 Detailed methodology for MICRON

#### Creating the most informative crop for training

For each image, we partitioned pixels into perceptually coherent regions using SLIC [45]. SLIC produces compact, uniform *superpixels* that respect object boundaries while substantially reducing the number of elements in the image to be processed. By default, we partitioned each image into 100 SLIC superpixels. For each superpixel, we defined a corresponding image crop as the bounding box enclosing the superpixel, with additional margins, and generated a binary mask. Shape irregularity of superpixels was quantified based on binary masks. We hypothesize that superpixels exhibiting greater deviation from a quadrilateral shape capture higher intra-region feature variation, are likely to contain more diverse cell-types, and are therefore more informative for downstream learning (Fig. S4). Based on this assumption, we selected the most *M* = 30 irregular superpixels and used their corresponding crops and binary masks for model training. More details of the crop definitions are provided in Fig. S1a.

The top 30 selected crops were ultimately used to generate a summarized, automated annotation of the image. We repeated the same crop-generating process across all images.

#### Multiple-instance learning with quantile-based aggregation

We used a fully convolutional network to generate spatially resolved predictions for each instance, followed by a quantile function that aggregates predictions across instances into a prediction for the entire sample (Fig. 1a). First, we defined a dataset *D*, comprised of a set of instances ℬ (see Table S1 for notation details). For a sample *i*, the collection of their instances is denoted as 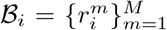 , where *m* specifies the *m*-th instance. The label (outcome) for the *m*-th instance of sample *i* is given as 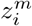, so that 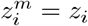 for all *m*.

Input images within a dataset, *D* all have dimensions of *h*× *h* ×*N*. After being processed by a fully convolutional network (FCN), the instance-level class probabilities for instance *m* in sample *i* are encoded in 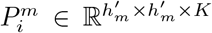. Here, 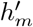 gives the number of pixels within instance *m*. These instance-level probabilities are subsequently aggregated by a quantile function 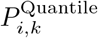, capturing the distribution of instance-level predictions for class *k* in sample *i* as,

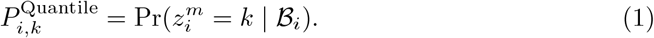

From this, we featurize distributions of predicted probabilities in each class with a quantile operation as,

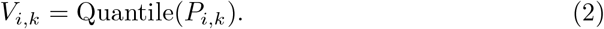

Computing this for each sample *i*, we obtain the quantiles of predicted probabilities for class *k*. In summary, quantile functions offer a discretized summary of the distribution of instance predictions to summarize each sample.

Each spatial location in this tensor 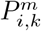 corresponds to an instance defined by the receptive field of the network, such that instance-level prediction scores for class *k* across the *M* instances can be defined as 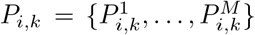. After sorting the values in *P*_*i,k*_ and filtering those instances that are not in the binary mask generated by SLIC, we obtain a filtered vector per class 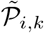. The values of 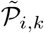 are ultimately used to compute a quantile-based representation. Specifically, given a predefined number of quantiles, *Q*, the *q*-th quantile is selected as the element at index

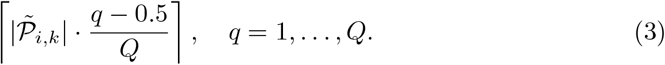

In this notation, · points to the index of the ordered values of 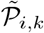, given as 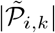 satisfying a particular quartile threshold, 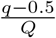.

Quantile vectors computed across all classes in sample *i* are concatenated into a joint vector **V**_*i*_, where **V**_*i*_ = [*V*_*i*,1_, … , *V*_*i,K*_]. In our implementation, we consider only binary outcomes, such that each **V**_*i*_ = [*V*_*i*,1_, *V*_*i*,2_]. Furthermore, the class 1 quantile vector in sample *i* is given as *V*_*i*,1_ = Quantile 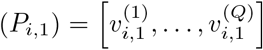 , such that each quantile value is computed according to equation (3) as,

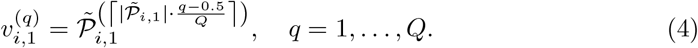

For each sample *i*, the concatenated quantile vector **V**_*i*_ is input to a softmax function, *S*_*i*_ = softmax(**V**_*i*_), to obtain the probability that the sample belongs to each class, 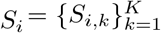. We then train the model using categorical cross-entropy loss over *I* total samples and *K* possible outcome labels as,

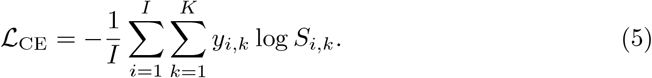

Here, *y*_*i,k*_ is a binary indicator encoding whether the ground truth label of sample *i* is outcome *k*. This loss function is used to optimize the learned representations for sample *i* by capturing concordance between true and predicted sample labels.

#### Scoring and prioritizing outcome-associated microenvironments

For each test sample, `MICRON` outputs an embedding vector for every crop. These embeddings are extracted from the second-to-last layer of the network and capture high-level feature representations. The *d*-dimensional embedding vector of the *c*-th crop in sample *i* is denoted as **X**_*i,c*_ ∈ ℝ^1*×d*^ (Fig. 1b). To stratify the contribution of each crop to the outcome prediction, we applied the SHAP TreeExplainer to a trained Random Forest classifier, using **X**_*i,c*_ as input [19]. For each crop *c* in sample *i*, SHAP assigns a contribution value to each of *d* embedding dimensions, where the SHAP value for each feature is denoted as,

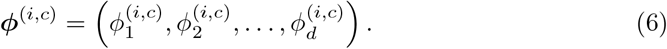

We then summarize the overall importance of crop *c* in sample *i* by computing the absolute sum of its SHAP values as,

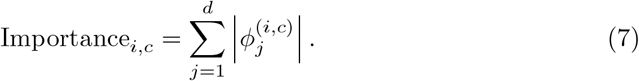

This score reflects the overall contribution of each crop embedding computed to the final prediction by aggregating the contributions across all embedding dimensions.

In all evaluation steps, we selected the top 20 highest-scoring crops in test images based on SHAP values and performed mean pooling over their embedding vectors to compute featurizations for each sample (Fig. 1b-c). This pooling step thereby creates sample featurizations, which reflect the most informative crops, while suppressing the less-relevant background information. We then conducted a second-stage evaluation in which the sample featurizations were used to train a random forest classifier to predict outcomes. This procedure was repeated over 200 random folds to improve robustness. For each fold, we recorded prediction probabilities for the test samples. The final AUROC and its standard deviation were obtained by averaging probabilities across all folds in the test set.

## 5 Supplementary information

Additional experimental results and details about `MICRON` and its evaluation are provided as Supplementary Information.

## 6 Availability of Data and Software

`MICRON` was evaluated on three publicly available datasets from Refs. [20] (Diabetes), [21] (brain tumors), and [6] (breast cancer). `MICRON` is available as open-source software at https://github.com/ChenCookie/MICRON.

## 7 Acknowledgments

This work was supported in part by grants R21AG084251 (to NS by the National Institute on Aging of the NIH), and R21AI171745 (to NS by the National Institute of Allergy and Infectious Diseases of the NIH).

## Supplementary Material

### 1. MICRON AND RELATED FEATURIZATION APPROACHES FOR IMC ANALYSIS

**Fig. S1.**
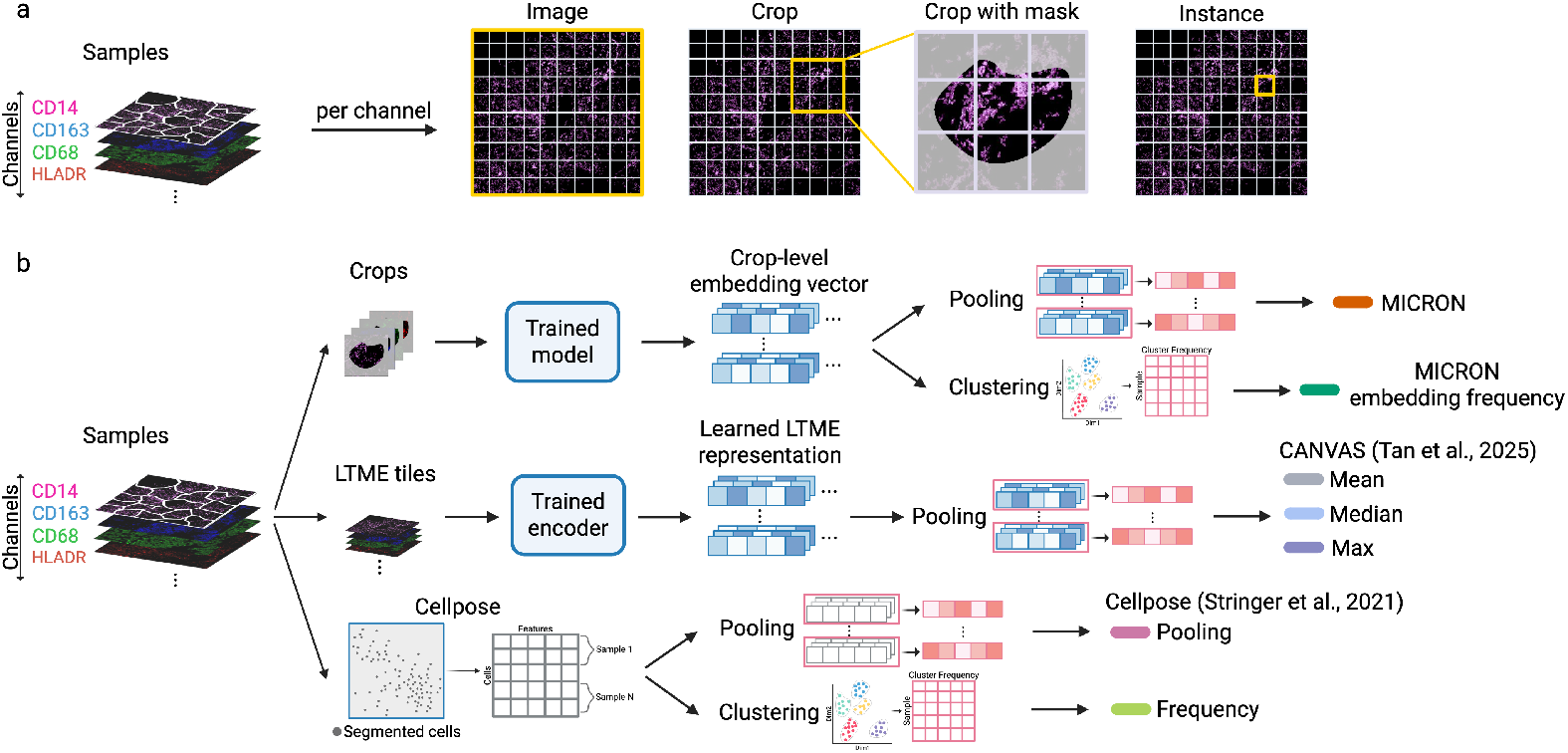
Overview of MICRON and comparison to Cellpose-based and CANVAS-based featurization approaches. (**a**) Illustration of how image regions are defined in `MICRON`. Multiplexed imaging samples consist of multiple channels. Each sample is cropped into several square regions, where each crop contains multiple instances. A bounding box binary mask is generated by SLIC and applied to define the region of interest within each crop. (**b**) Comparison of `MICRON` with tile-based and segmentation-based approaches. In `MICRON`, each crop-level embedding vector is generated by a trained model. These embeddings are aggregated either by pooling to produce a sample-level representation or by clustering to compute embedding frequency features. In the CANVAS framework, local microenvironment (LTME) tiles are extracted and encoded using a trained encoder. Sample-level representations are obtained via global pooling operations (mean, median, or max) across tile embeddings. To define features via CellPose segmentations, the expression of each protein can either be pooled over all segmented cells or cells can be clustered to define frequency features.

### 2. ADDITIONAL BIOLOGICAL INSIGHTS GLEANED ACROSS DATASETS

**Fig. S2.**
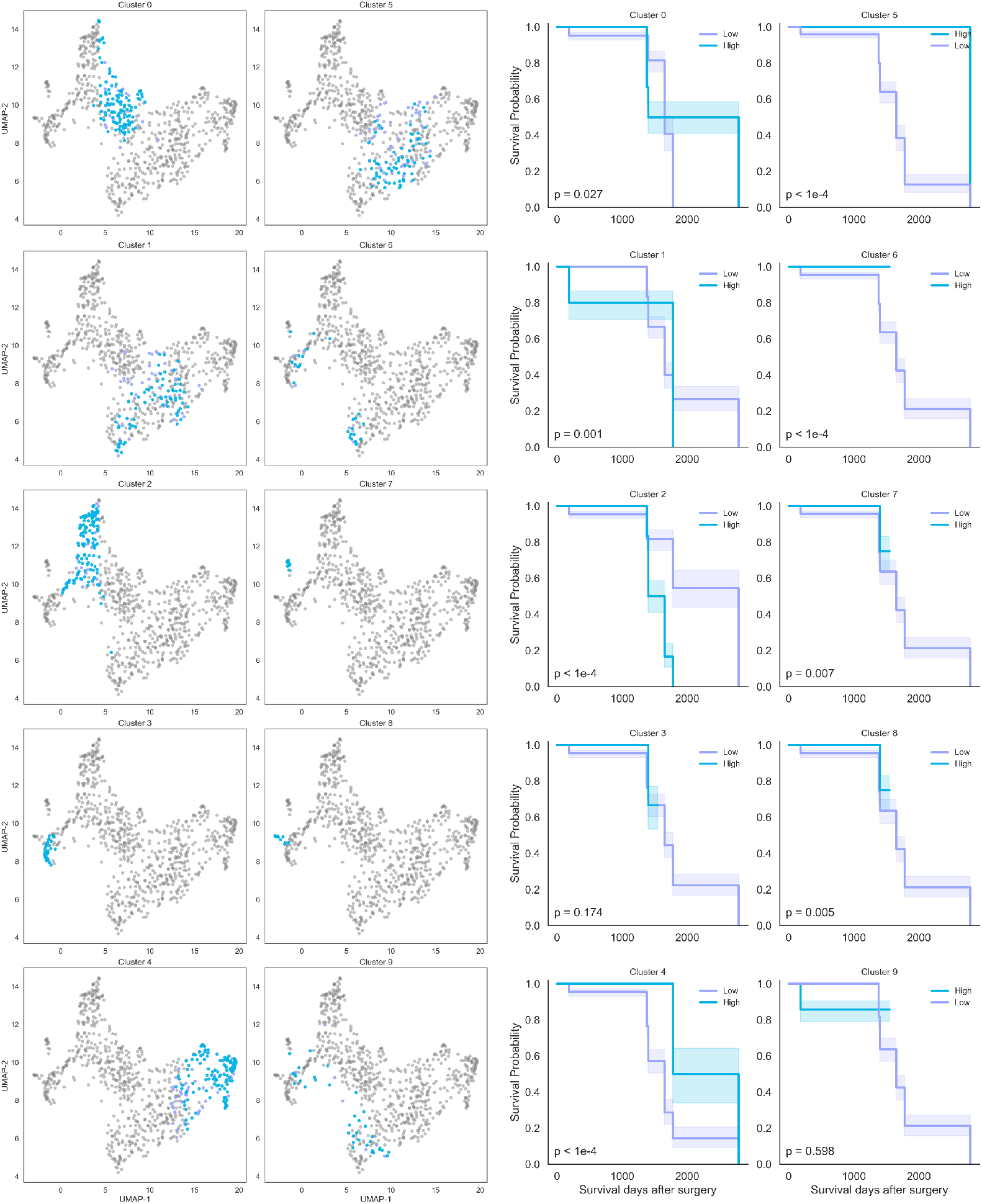
Identifying survival-associated microenvironments. To evaluate the prognostic relevance of the learned superpixel representations under `MICRON`, we clustered superpixels across samples using *k*-means (*K* = 10) and computed the frequency of each cluster per patient. For a given cluster, patients were stratified into high- and low-frequency groups based on the 75th percentile frequency of that particular cluster. The UMAP visualizations provide two-dimensional projections of superpixels identified across images. Blue points denote cells belonging to the indicated cluster. Kaplan–Meier survival analysis was used to quantify the differences in survival between patients with high and low frequencies of a given cluster. Several clusters showed significant survival differences between high and low frequency groups. Cluster 4 showed the most significant association with survival (*p <* 10^−4^).

**Fig. S3.**
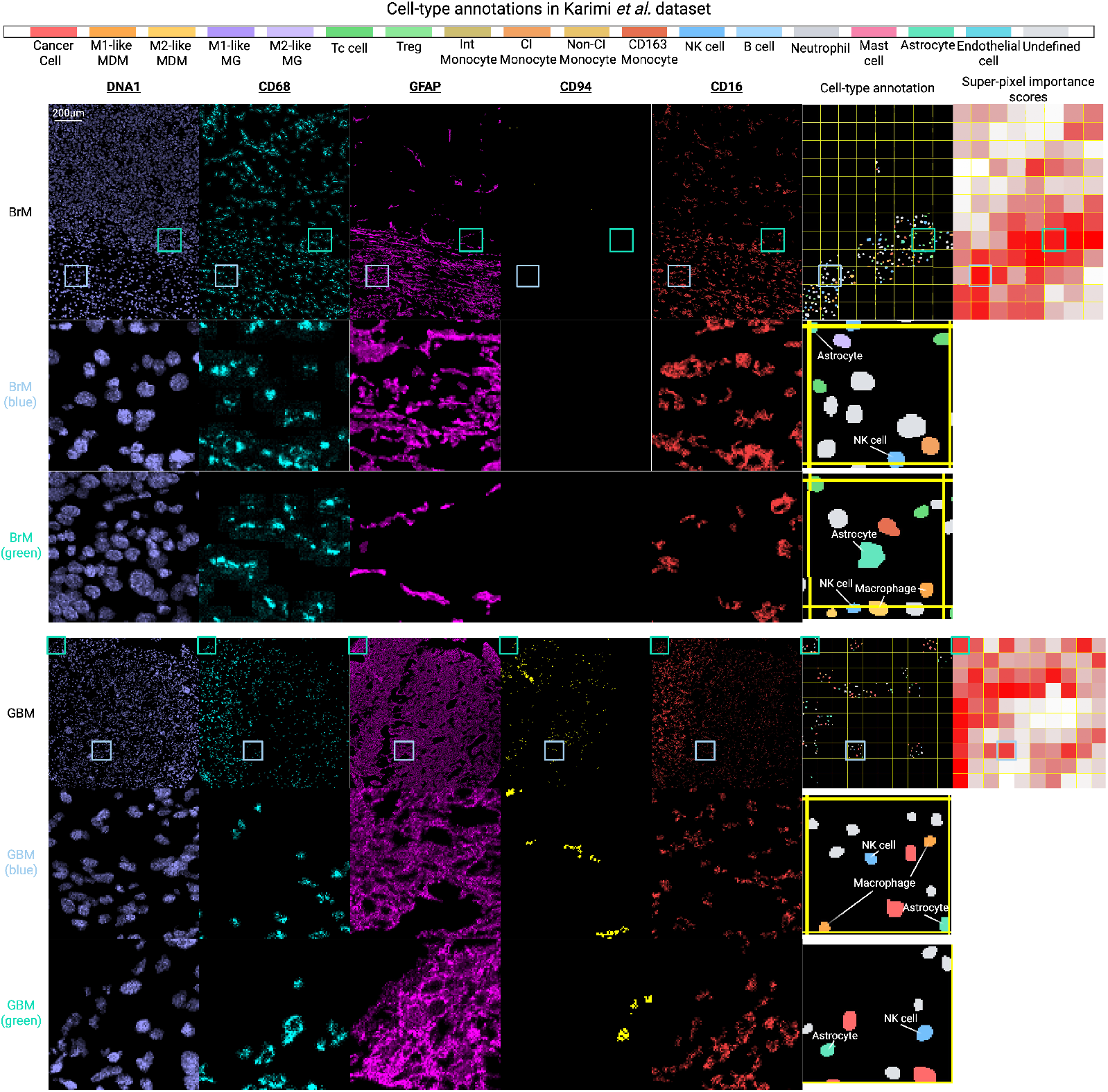
Visualization of high-importance superpixel regions and associated cell-type markers in the Karimi brain tumors dataset. Representative superpixel regions with high importance scores identified by `MICRON` are shown for brain metastasis (BrM) and glioblastoma (GBM) samples (heatmaps on right). Images show expression for individual channels, including DNA1, CD68, GFAP, CD94, and CD16. DNA1 used to identify nuclei and determine cellular localization. Canonical markers were used to define key cell types, including *CD*68^+^ macrophages, *GFAP*^+^ astrocytes, and *CD*94^+^*CD*16^+^ NK cells. Blue and green squares denote two different superpixels selected and deemed to be outcome-associated. Corresponding celltype annotations illustrate the spatial distribution of annotated cell populations. Zoomed-in views reveal spatial proximity among astrocytes, macrophages, and NK cells within the selected regions. Although CD94 expression was relatively low in BrM samples, *CD*94^+^ signals were occasionally present but visually subtle in the corresponding images.

**Fig. S4.**
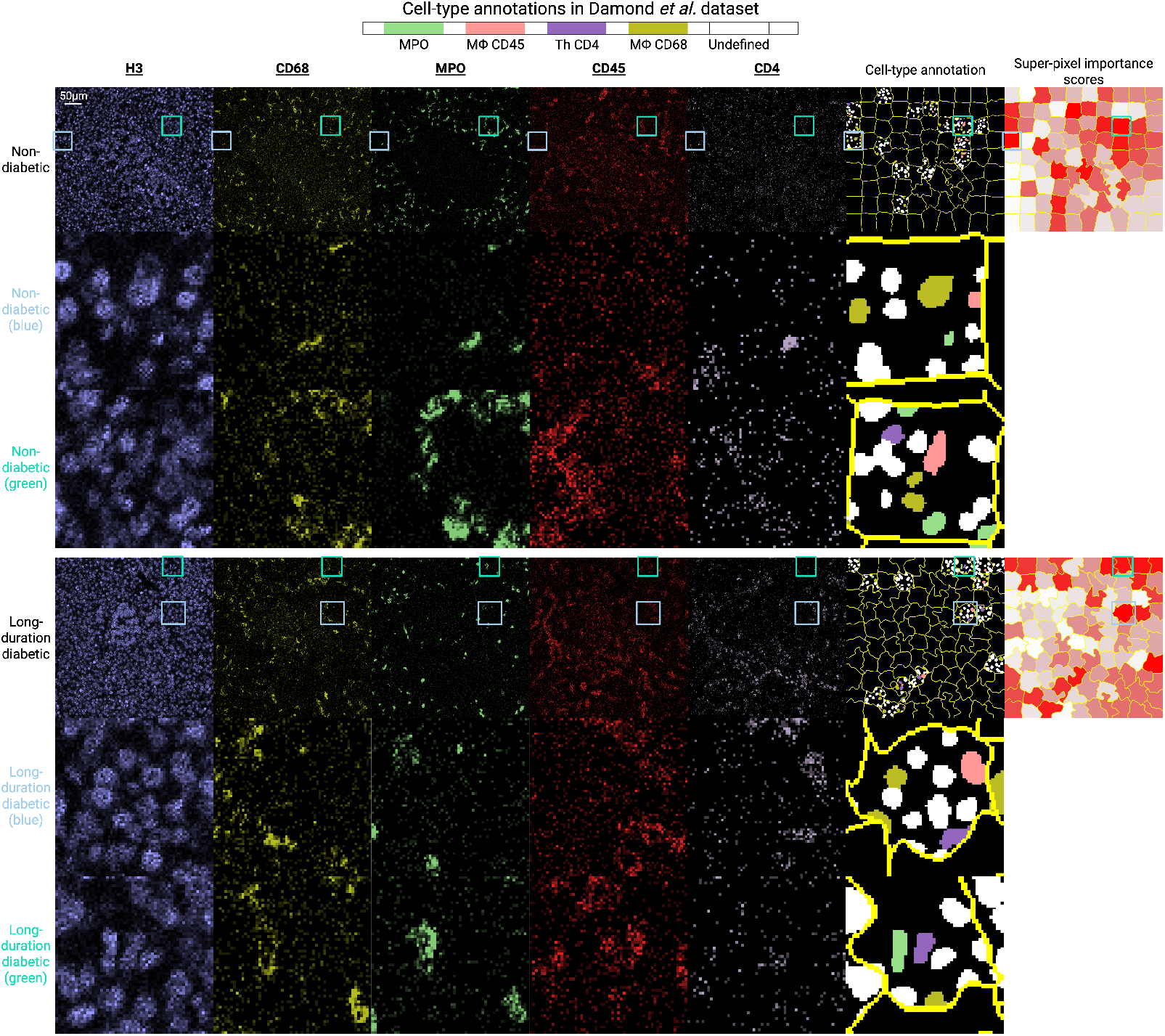
Visualization of high-importance superpixel regions and associated immune cell markers in the Damond et al. diabetes dataset. Examples of superpixel regions with high importance scores predicted by `MICRON` are shown for non-diabetic and long-duration diabetic samples. Images show expression for individual markers, including H3, CD68, MPO, CD45, and CD4. H3 staining was used to identify cell nuclei. Immune cell populations were annotated based on canonical markers, including *CD*68^+^ macrophages, *MPO*^+^ neutrophils, *CD*45^+^ leukocytes, and *CD*4^+^ T helper cells. Light-green and light-blue boxes mark two representative superpixels with high model importance that were prioritized as being outcome-associated by the `MICRON` framework. The cell-type annotation maps display the spatial distribution of immune cells in these regions. The samples were not arcsinh-transformed during preprocessing, which causes the cropped regions to appear more irregular in shape. Enlarged views highlight the presence and local organization of immune cell populations within the influential superpixel areas identified by `MICRON`.

**Table S1.**
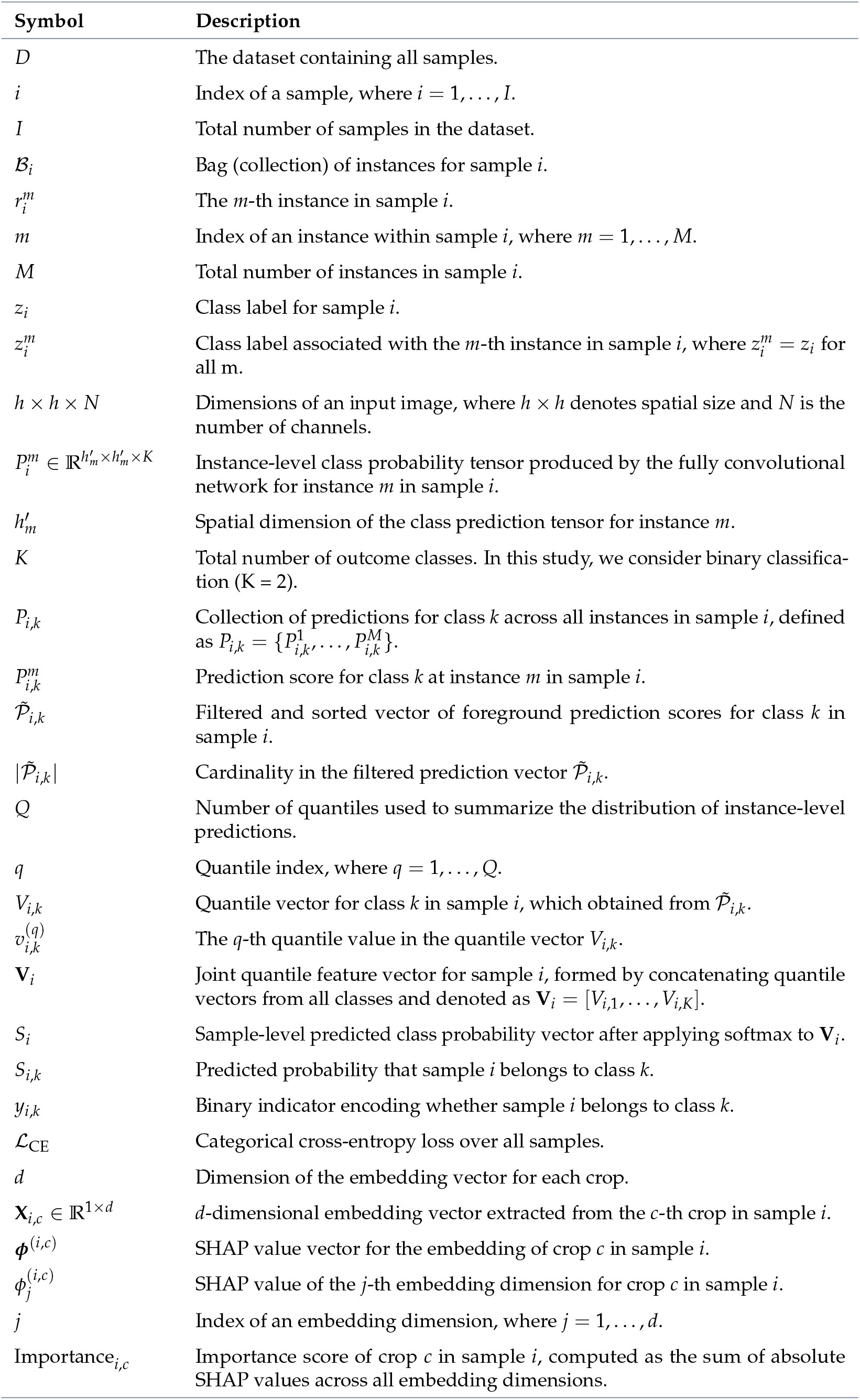
Notation used in MICRON.

